# Stator pool size shapes the mechanosensitivity of the bacterial flagellar motor

**DOI:** 10.64898/2026.07.27.740596

**Authors:** Brennen M. Wise, Shabduli A. Sawant, Navish Wadhwa

## Abstract

The bacterial flagellar motor adapts to mechanical load by adding or removing stator units, which can either be inactive in the inner membrane (the stator pool) or attached to the motor in a torque-generating configuration. This adaptation is well understood, but what sets the load range over which the motor responds remains unclear. Building on a previous two-state model of stator binding, we derive how the on-rate depends on the diffusing stator pool and find that it increases in proportion to pool size. The model predicts that pool size governs motor occupancy at low load but not at high load, where the motor saturates regardless of how many stators are available. We confirmed this prediction in *Escherichia coli*: raising stator expression increased swimming speed and single-motor rotation rate at low load. This dependence on expression weakened as load rose and was undetectable at high load. The model further predicts that larger pools make the motor mechanosensitive at lower loads, which our measurements of load-induced occupancy change support. Mechanosensitivity is therefore not fixed by motor architecture alone, but also determined by stator abundance, which cells can set through expression to position their mechanosensory range.

**Significance:** Using a statistical mechanics model of stator binding, we predict how flagellar motor output depends on mechanical load and on the number of stators a cell expresses. Combining the model with measurements of swimming cells and single motors, we find that the size of the stator pool sets the range of load over which the motor is mechanosensitive. Mechanosensitivity is therefore a property the cell can tune through stator expression, not a fixed feature of the motor, a principle that may extend to other force-responsive molecular complexes assembled from a limited pool of components.

## Introduction

To find resources, escape harm, or infect a host, bacteria must navigate both the chemical and the physical features of their environment [1]. Many species, including the model organism *Escherichia coli*, do this with the flagellar motor, a rotary molecular complex that spans the cell envelope and turns an extracellular filament, the flagellum, to power swimming [2–4]. Switching the motor’s rotation between counterclockwise and clockwise alternates straight “runs” with reorienting “tumbles,” letting the cell navigate its chemical environment by chemotaxis [5–7]. In many species, the motor also responds to physical inputs: it raises its torque output when the mechanical load on the flagellum rises and lowers it when the load falls [8–10]. This mechanosensitivity has generally been treated as a fixed property of the motor, set by its architecture.

The torque that drives rotation, and the mechanosensitive response itself, both come from the stator units. These protein complexes harness an electrochemical gradient of cations—H^+^ ions in *E. coli*—to power flagellar rotation [11–14]. Each stator consists of two proteins, MotA and MotB, assembled in a 5:2 stoichiometry [15, 16]. MotB anchors in the periplasm, while MotA exerts torque on the flagellar rotor via electrostatic interactions with the rotor protein FliG [17, 18]. Structural and recent *in vivo* evidence indicates that MotAB generates torque through a rotational mechanism, in which the MotA pentamer rotates around the MotB dimer [19, 20].

Despite their name, stators are not static. They exchange continually between the motor and a reservoir of inactive units in the inner membrane, the stator pool (Fig. 1A) [21]. Because each bound stator contributes roughly equal torque, the motor’s rotation rate reports the number of bound stators [22–24]. This has enabled bead and tethered-cell assays, often combined with load manipulation by electrorotation, viscous agents, or magnetic tweezers, to track stator number as load is varied [24–26]. These experiments revealed that the stator complex is mechanosensitive: the number of stators bound to a motor changes with mechanical load [8, 9]. The proposed mechanism centers on a catch bond between the stator and the peptidoglycan layer [26], with recent evidence indicating that long-range interactions and flexible domains regulate this bond [27]. As motor load increases, this bond strengthens, prolonging stator dwell time and thus increasing occupancy [24, 28–30]. In this picture, mechanosensitivity arises from the load-dependent off-rate: load determines how readily bound stators leave.

**Figure 1.**
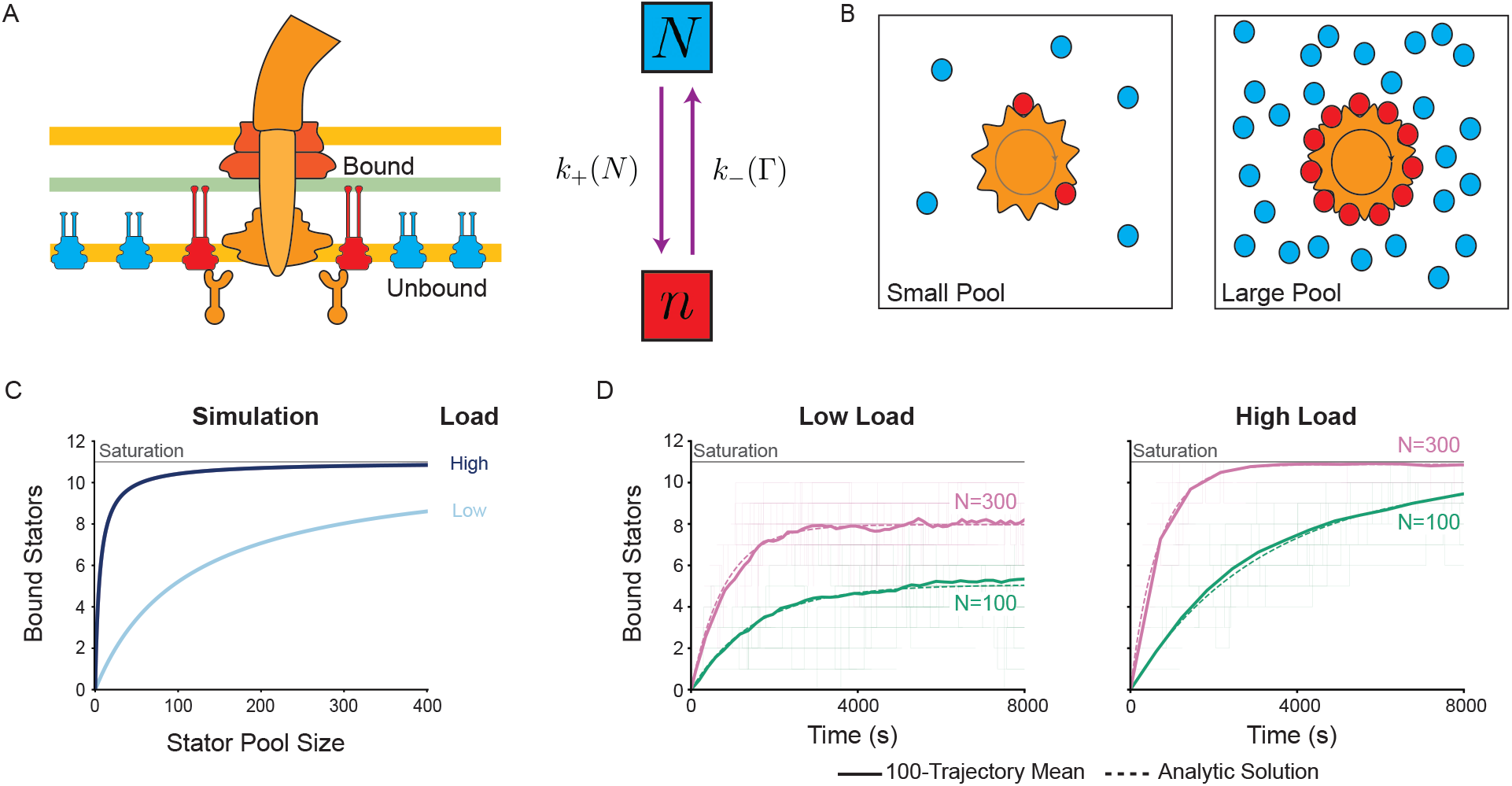
Stator occupancy at the flagellar motor is tuned by load and stator pool size. A) Schematic of the flagellar motor and the two-state model with bound stators in red, unbound stators in blue, and the peptidoglycan layer in green. Stators are partitioned into two states: bound (*n*) and unbound (*N* ). The off-rate depends on motor torque (Γ) and the on-rate depends on the number of stators in the pool (*N* ). **B)** Cross-section of a rotor (orange), with bound and unbound stators shown. As the number of stators in the pool increases, we expect that the number bound to the motor will also increase, eventually reaching saturation. **C)** Steady-state stator occupancy (*n*_ss_) as a function of stator pool size (*N* ), computed from Eq. (3) for a range of torque energies (*ϵ*_*T*_ ) spanning low to high load. **D)** Collection of individual stochastic simulations (thin lines), average of 100 simulations (thick line), and analytical result (dashed line). Green is a low stator pool size (*N* = 100) and pink a large one (*N* = 300). The left panel shows low load, the right panel high load.

What governs the other side of this balance, the rate at which stators bind, has received less attention. A cell maintains a pool of roughly 100 stators [21], yet a single motor holds at most 11 [31] and the motor of a freely swimming cell operates with only 5 to 6 [32]. The pool therefore far exceeds what any one motor can use. Raising stator expression increases the number of stators engaged at the motor [33–35], but whether the pool that supplies them sets the motor’s mechanosensitivity is unknown. More generally, the abundance of a component that exchanges with a pool could set where on the load axis a force-responsive complex becomes sensitive [36, 37], a possibility that can be tested in the well-characterized flagellar motor.

Here, we extend the two-state model of stator binding by deriving how the on-rate depends on the pool, and we find that it increases in proportion to pool size. The model, parameterized from prior measurements, predicts that pool size governs stator occupancy at low load but not at high load, where the motor saturates regardless of how many stators are available. We test this in *E. coli*, first in populations of swimming cells across a range of loads, where swimming speed rises with stator expression at low load and converges at high load, and then at the level of single motors, where tethered-cell rotation shows the same load-dependent pool effect. The same model predicts that pool size sets the load range over which the motor is mechanosensitive, with larger pools becoming sensitive at lower loads, and measurements of load-induced occupancy change support it. Thus, a cell can position the load at which its motors respond by how many stators it expresses, making mechanosensitivity a tunable cellular parameter rather than a built-in feature of the motor.

## Materials and Methods

### Gillespie Algorithm

To simulate stator remodeling, we used the Gillespie algorithm, a stochastic method for simulating reaction kinetics [38]. We initialized each simulation by setting the pool size (*N* ) and the number of bound stators (*n*). The total reaction rate was

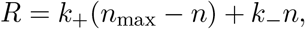

where *n*_max_ = 11 is the maximum stator occupancy and *k*_+_ and *k*_−_ are the stator binding and unbinding rates. We drew the time until the next reaction (*τ* ) from

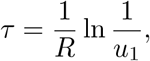

where *u*_1_ ∼ Uniform(0, 1). The probability that the next reaction was a binding event was

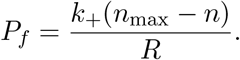

We then drew *u*_2_ ∼ Uniform(0, 1): if *u*_2_ *< P*_*f*_, the reaction was a binding event; otherwise it was an unbinding event. We repeated this process until 40 reactions had occurred.

### Parameters for Simulations

To set the timescale of our Gillespie simulations, we included a base rate *k*_0_, estimated from Shi et al., Tipping et al., and Wadhwa et al. [9, 24, 33]. We estimated the bound energy *ϵ*_*b*_ and torque energy *ϵ*_*T*_ from Wadhwa et al. [24], and chose the unbound energy *ϵ*_sol_ to be smaller than both. Specifically, the torque energies for tethered cells in different load conditions reported in Wadhwa et al. were on the order of 1-10 *k*_*B*_*T* [24]. All energies are reported in units of *k*_*B*_*T* . To estimate *γ*, the ratio of a stator’s cross-sectional area to the total inner-membrane area, we modeled the cell as a cylinder of length 1 *µ*m and radius 0.35 *µ*m, giving an inner-membrane surface area of approximately 3 *µ*m^2^. We took the diameter of a stator from previous work and doubled it to account for crowding effects [15]. Following Leake et al., who measured an average of 200 MotB molecules per cell, we used a baseline pool of *N* = 100 diffusing stators (2 motB per stator) [21]. Finally, we set *n*_max_ = 11 [31].

### Bacterial Strains and Cultures

We used *E. coli* strain MG1655 as the wild-type for all experiments. To tune stator expression, we constructed a strain in which the chromosomal *motAB* genes were deleted and stators were expressed from the arabinose-inducible plasmid pBAD33 [NW166 = MG1655 Δ*motAB* + pBAD33(*motAB*)]. For tethered-cell assays, we used strain NW263 [NW166 + pFD313(fliCst)], which expresses a sticky flagellin variant that lets individual motors be tethered to a glass surface. For Western blots, we added a 6×His-tag to the C-terminus of MotB in the *motAB*-pBAD33 construct using Agilent’s QuikChange II site-directed mutagenesis kit, and transformed the resulting construct into *E. coli* BL21 for protein expression.

Cultures were grown overnight at 37^◦^C in LB medium supplemented with 25 *µ*g/mL chloramphenicol for NW166 and with 25 *µ*g/mL chloramphenicol plus 100 *µ*g/mL ampicillin for NW263. We diluted 100 *µ*L of overnight culture into 10 mL of terrific broth (TB) with the appropriate antibiotic and grew it at 37^◦^C with shaking to an OD_600_ of 0.5. We then added arabinose to a final concentration of 0.002, 0.02, 0.2, or 2% w/v and grew the cultures for an additional 30 min. All assays were performed immediately after growth with arabinose.

### Western Blot

Strains were grown in LB supplemented with 25 *µ*g/mL chloramphenicol to an OD_600_ of 0.4, induced with arabinose (0, 0.002, 0.02, 0.2, and 2%), and grown to a final OD_600_ of approximately 1.0. Cells were harvested by centrifuging 1 mL of culture at 4500×*g* for 5 min and lysed in lysis buffer (17.2 mM Tris-HCl [pH 7], 8.6 mM EDTA [pH 8], and 1 mg/mL lysozyme) at 37^◦^C and 500 rpm for 1 h. We mixed 60 *µ*L of the lysate with 100 *µ*L of 1× Laemmli buffer (60 mM Tris-HCl [pH 7], 10% glycerol, 2% SDS, 0.05% bromophenol blue) and boiled it at 95^◦^C for 10 min. We loaded 20 *µ*L of lysate into SurePAGE precast polyacrylamide gels (Genscript), resolved them for 1 h at 120 V, and transferred them onto a PVDF membrane with the iBlot 2 gel transfer device at 25 V for 6 min. Membranes were probed with a 1:10000 dilution of anti-His-tag antibody (Invitrogen [MA1-135]) followed by a 1:1000 dilution of horseradish peroxidase-conjugated goat anti-mouse immunoglobulin G (Invitrogen [31430]). Blots were developed with Pierce ECL substrate (Thermo Fisher Scientific) and visualized on an Azure Biosystems imager in the chemiluminescence channel with 3×3 binning and a 20 s exposure.

### Swimming Speed Measurements

Cell cultures (1 mL) were pelleted by centrifugation (1500 × *g*, 7 min) and resuspended in 1 mL of motility buffer (MB; 66.7 mM NaCl, 0.1 mM EDTA, 10 mM K_2_HPO_4_), supplemented with 10% or 15% Ficoll 400 (Bio Basic) where indicated. Cells remained in the indicated buffer for 5−10 min before imaging. Imaging chambers were prepared by attaching a GeneFrame (Thermo Fisher) to a microscope slide, adding 100 *µ*L of cell suspension, and sealing with a glass coverslip. We imaged cells by phase-contrast microscopy (20× objective) for 1 min at 20 fps on a Nikon Optiphot-2 with a ThorLabs DCC1545M camera. We tracked cell positions across these recordings and computed average single-cell swimming speeds from the trajectories using custom scripts in Python and ImageJ. As expected, swimming speed varied within each population, so we used the population mean as a summary statistic for each condition. Experiments were performed with several biological replicates (n = 6, n = 5 (WT) for 0% Ficoll; n = 3 for 10% and 15% Ficoll for all strains). An additional biological replicate was performed to test if the speed of cells increased beyond 2% arabinose at 0% Ficoll. This one replicate (found in the supplement) used the same MB except supplemented with 1 mM methionine.

### Tethered-Cell Assays

Cell cultures (5 mL) were pelleted by centrifugation (1500 × *g*, 7 min) and resuspended in 1 mL of motility buffer. Flagella were sheared by passing the cell suspension through 7 cm of tubing (0.023” inner diameter, Science Commodities Inc.) 150 times between two syringes (3 mL syringe and 23 ga Luer stubs, Fisher Scientific). Cells were pelleted again (3000 × *g*, 3.5 min) and resuspended in 1 mL of motility buffer. The suspension was added to a tunnel slide (a channel formed by a glass slide and coverslip separated by two strips of double-sided tape), which was left inverted for 10 min to promote surface attachment. Unattached cells were removed by washing twice with 200 *µ*L of MB, and 200 *µ*L of the appropriate buffer (MB with 0%, 10%, or 15% Ficoll) was then flowed through the slide. For each condition, at least 5 fields of view of 1000 × 1000 pixels (≈1800 *µ*m^2^) were imaged for 30 s at 50 fps on a Nikon Eclipse Si with a FLIR Blackfly S camera using a 40× objective. Object detection, rotation-frequency calculation, and data analysis were performed with custom scripts in Python. Experiments were performed with several biological replicates (n = 3 for each Ficoll condition).

### Statistical Tests and Samples

Statistical comparisons were performed with TOST (p-values denoted *p*_*T*_ ) and one-way ANOVA (p-values denoted *p*_*A*_), implemented with SciPy and statsmodels in Python [39, 40]. We used TOST to determine if two groups are essentially identical, and one-way ANOVA to test for differences across groups, with a significance level of *α* = 0.05 for all tests. The equivalence interval for TOST was determined before tests were performed. We chose our speed interval to be approximately 33% of the smallest mean speed in all of our data at 0% Ficoll (9 *µ*m/s *→ ±*3 *µ*m/s for swimming cell assays) and we chose our frequency interval to match the approximate frequency contribution of a single stator in a tethered cell, being 1.3 Hz [24]. All experiments included biological replicates (Tables S2, S3, S4). Statistics were calculated based on replicate means. Summary tables of statistical results can be found in the supplement (Tables S5, S6).

## Results

### Pool size sets the stator binding rate

Our model extends an experimentally grounded, two-state description of stator dynamics, in which a stator occupies either a diffusing state in the membrane or a bound state at the motor, with transitions set by the binding and unbinding rates (Fig. 1A) [24, 26, 41]. The average number of bound stators evolves as:

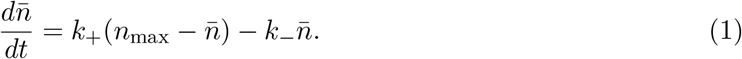

Here, 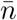 is the average number of bound stators, *n*_max_ is the maximum occupancy, and *k*_+_ and *k*_−_ are the binding and unbinding rate constants. Their ratio is given by 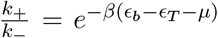, where *β* = 1*/k*_*B*_*T* and *k*_*B*_*T* is the thermal energy. The energy of a bound stator is *ϵ*_*b*_ − *ϵ*_*T*_, where *ϵ*_*b*_ is the bound-state energy and −*ϵ*_*T*_ is the contribution from torque generation. As load increases, the torque per stator rises, raising *ϵ*_*T*_ and making binding more favorable. The final term, *µ*, is the chemical potential for a stator moving from the diffusing to the bound state [24].

Earlier models treated *µ* as a constant. We instead derive it from the free energy of the diffusing pool (see Supporting Material for details). Our approach uses a lattice model where we assume the inner membrane can be divided into evenly sized lattice sites, Ω, each of which can hold a single stator. For a more physical interpretation, we define 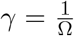, where *γ* represents the ratio of a stator’s cross-sectional area to the total inner-membrane area, a measure of how many stators the membrane can hold. The final expression for the chemical potential is

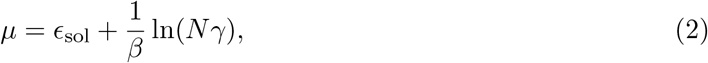

where *N* is the number of stators in the pool. We assign the load dependence entirely to the off-rate through the stator–peptidoglycan catch bond, consistent with prior work [24, 26], so that pool size enters only through the on-rate. Because *µ* enters *k*_+_, the binding rate then grows with pool size such that *k*_+_ ∝ *N* (Fig. 1B): the more stators in the pool, the faster a motor recruits them. This prediction is consistent with single-motor measurements in which the slow component of stator turnover speeds up at higher stator expression, as expected if it depends on the size of the membrane pool [33]. Solving Eq. (1) at steady state gives the occupancy as:

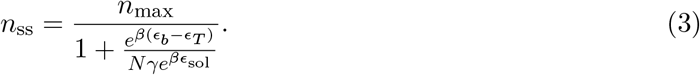

Evaluating this expression with parameters constrained by prior measurements (Table S1) predicts the number of bound stators across expression levels and loads (Fig. 1C): steady-state occupancy rises with both the torque per stator and the pool size *N* .

### Pool size effects on motor occupancy depend on load

To explore these predictions, we simulated stator binding and unbinding with the Gillespie algorithm (see Methods), using the rate constants above. Each trajectory follows the number of bound stators as it approaches steady state. As expected, a larger pool reaches a higher steady-state occupancy and reaches it faster. Averaging 100 trajectories recovers the analytical solution of Eq. (1) (Fig. 1D).

We then repeated the simulations at higher load, modeled by increasing *ϵ*_*T*_ to reflect the greater torque each stator generates against a larger external load. At high load, motors approach maximum occupancy regardless of pool size; only the time to reach saturation depends on *N* (Fig. 1D). Pool size therefore sets steady-state occupancy at low load but not at high load, where it sets only the kinetics. Because occupancy determines torque output, the model predicts that cells with different pool sizes swim at different speeds at low load but at the same speed at high load, differing only transiently as occupancy approaches steady state.

### Pool size effects on swimming motility change with load

The model predicts that cells with larger stator pools recruit more stators to the motor. At low Reynolds number, swimming speed is proportional to flagellar rotation rate, so a higher stator occupancy should make cells swim faster. To test this prediction *in vivo*, we expressed *motAB* from the arabinose-inducible plasmid pBAD33 [42–44]. Raising the arabinose concentration increased MotB levels, indicating a larger stator pool because MotA and MotB are coexpressed from the same construct (Figs. 2A, S1). We tracked freely swimming populations under a light microscope to measure their speeds (Fig. 2B,C). To set a threshold for excluding the non-motile cells occasionally present in the field, we measured Δ*motAB* cells across several populations and found an average speed of ≈4 *µ*m/s from Brownian motion. Since the speeds of the fastest of these overlapped with those of the slowest motile cells, we applied a 4 *µ*m/s lower threshold to exclude most non-motile cells from the dataset, which did not affect the overall trend (Fig. S2A). In motility buffer (MB), populations expressing more stators swam faster than those expressing fewer (Fig. 2D). One-way ANOVA confirmed that swimming speed varied with stator expression (*p*_*A*_ *<* 0.001). Wild-type cells (native *motAB*, without the pBAD33 plasmid) showed little change in motility with arabinose at any load (*p*_*A*_ *>* 0.17), confirming that stator expression drove the speed differences, not the arabinose itself (Fig. S2B, Table S3).

**Figure 2.**
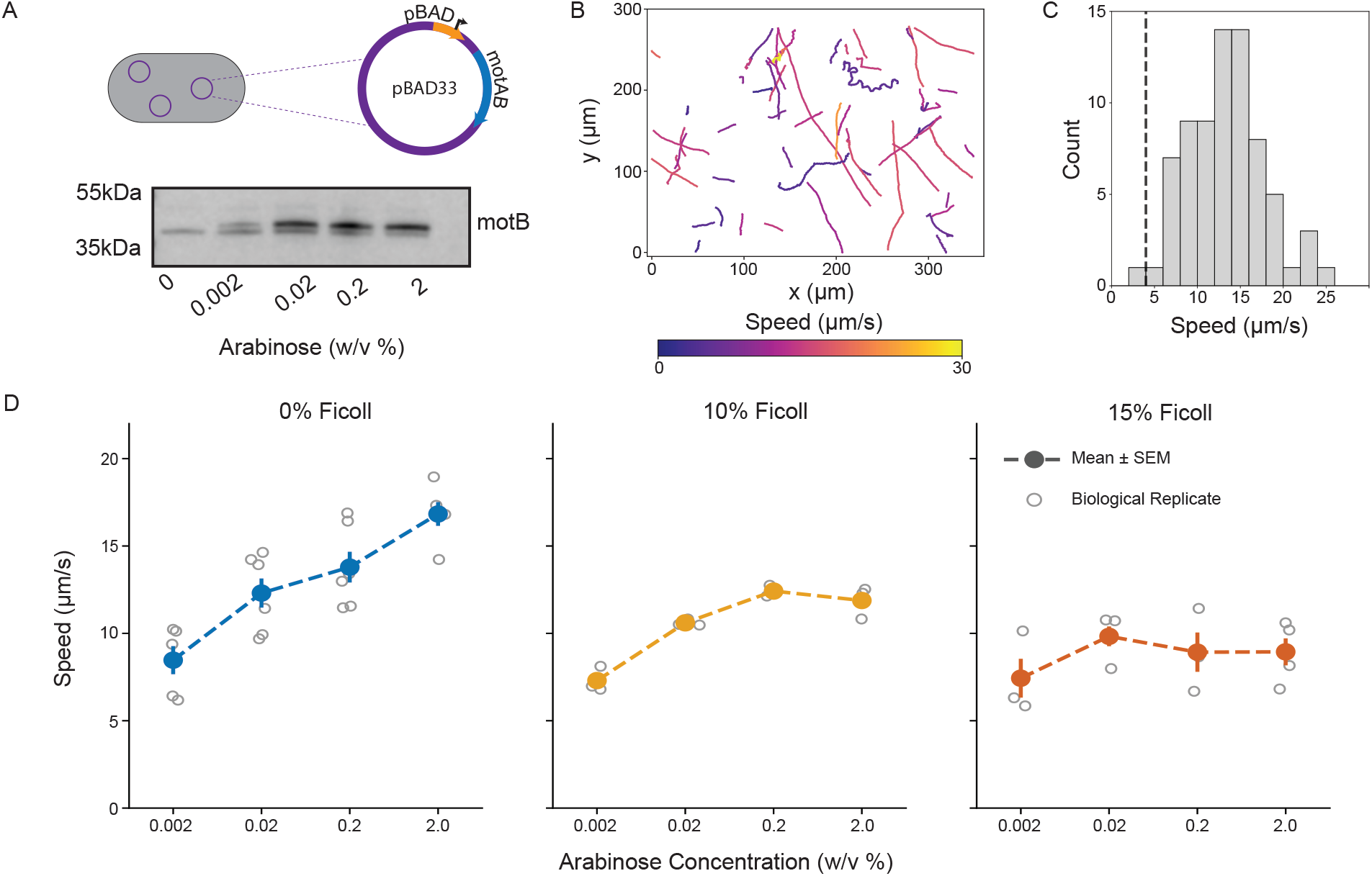
Swimming speeds of cell populations increase with stator expression at low load but not at high load. **A)** (Top) Schematic of the pBAD33 plasmid with an arabinose-inducible promoter and the inserted *motAB* gene. (Bottom) Western blot with 6×His-tagged MotB at different arabinose concentrations. Protein ladder markers are shown to the left. The full blot is shown in Fig. S1. **B)** Tracks of cells as detected by TrackMate. Color indicates the average speed of each track. **C)** Distribution of swimming speeds for a single population of cells. The dashed line denotes the non-motile speed cutoff (*<* 4 *µ*m/s). **D)** Average swimming speeds of cell populations at various stator expression levels in 0%, 10%, and 15% Ficoll (blue, yellow, and orange respectively). The error bars represent one standard error above and below the mean, and gray circles are the averages of individual biological replicates.

We then added Ficoll to MB as a viscous agent, raising the load on every motor and lowering the stator off-rate. At 10% Ficoll the trend persisted (*p*_*A*_ *<* 0.001), except that the two highest expression levels now appeared to swim at the same speed. An equivalence test (TOST) confirmed this: their speeds were statistically equivalent (*p*_*T*_ = 0.017). At 15% Ficoll, ANOVA detected no significant difference among populations (*p*_*A*_ = 0.478, Fig. 2D), but TOST did not establish equivalence either, leaving the 15% data inconclusive (Table S6). Even so, the overall pattern, from a strong dependence on expression at 0% Ficoll to none detectable at 15%, suggests that the effect of stator pool size on swimming speed weakens as external load increases. Our model’s calculated steady-state occupancy qualitatively reproduces this trend (Fig. 1C).

This qualitative load dependence follows directly from the model. At high load the off-rate approaches zero, so stators accumulate until every binding site is occupied regardless of *N* . Eq. (3) captures this, giving *n*_ss_ = *n*_max_ for large *ϵ*_*T*_ . At intermediate load, occupancy depends on *N* : high-expressing cells saturate their motors while low-expressing cells equilibrate below saturation. This matches the convergence of the two highest expression levels at 10% Ficoll (Fig. 2D).

### Pool size effects on tethered cells change with load

To confirm that this effect acts at the level of a single motor, we measured the rotation of individual motors in tethered cells, where rotation rate reports stator occupancy [22–24]. We transformed NW166 with pFD313, which carries a sticky filament gene [45], sheared the flagella, and attached cells to a glass coverslip by their filament stubs so that a single motor rotated the cell body (Fig. 3A). We extracted rotation frequency from phase-contrast trajectories recorded at 50 frames per second (Fig. 3B). To limit sampling bias, we recorded several fields for 30 s each and analyzed every rotating cell our detection method could track. Since rotation frequency decreases with cell size, we also applied a consistent cell-area threshold across conditions.

**Figure 3.**
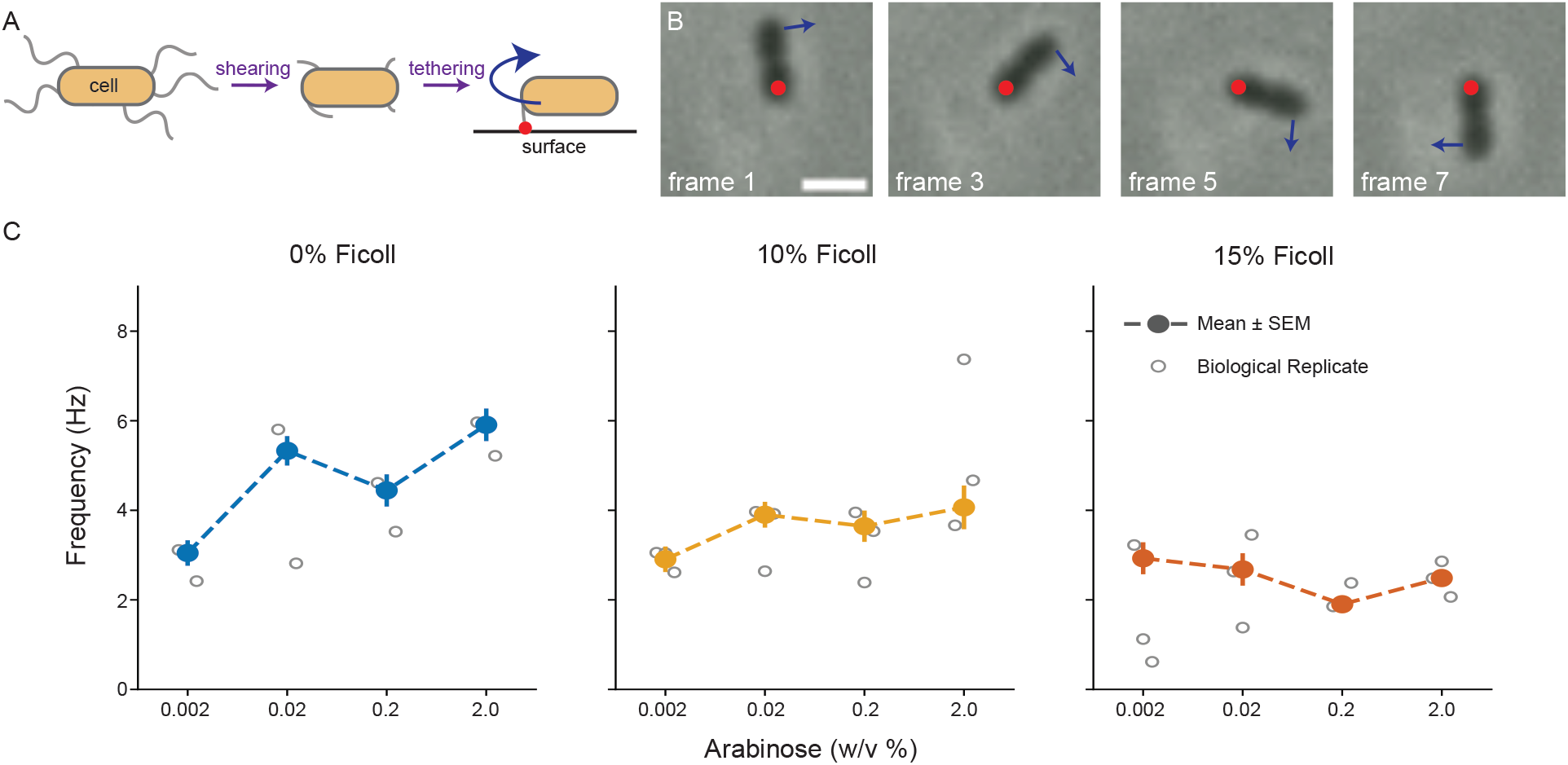
Single-motor rotation increases with stator expression at low load but not at high load. **A)** Flagellar filament stubs are attached to a glass coverslip in the tethered-cell assay. Rotation of the cell body is shown in blue and the attachment point in red. **B)** Raw phase-contrast images of a rotating cell under a 40× objective. The blue arrow shows the direction of rotation and the red dot marks the attachment point. The scale bar represents 1.5 *µ*m. **C)** Average rotational frequency of tethered cells at various stator expression levels in 0%, 10%, and 15% Ficoll (blue, yellow, and orange respectively). The error bars represent one standard error above and below the mean, and gray circles are the averages of individual biological replicates.

Without Ficoll, motors rotated faster as stator expression increased (*p*_*A*_ *<* 0.001, Fig. 3C). At 10% and 15% Ficoll, the change in frequency appeared minimal (Fig. 3C). ANOVA found no significant differences (*p*_*A*_ = 0.149 and 0.091 for 10% and 15% Ficoll), and TOST showed that the frequencies of cells in 15% Ficoll at the two highest arabinose concentrations were equivalent (*p*_*T*_ = 0.023, Table S6). To check that the motor’s directional bias did not shift across conditions, we measured each cell’s counterclockwise (CCW) bias; the mean did not differ between conditions (Fig. S3). Single motors thus reproduce the load-dependent pool effect seen in swimming assays, indicating that it is present at the level of a single motor and not a consequence of population-level or hydrodynamic effects in

### Stator pool size affects mechanosensing

To quantify mechanosensitivity, we wrote stator occupancy as *r* = *n*_ss_*/n*_max_, with 0 *< r <* 1 and *r* = 1 a fully occupied motor. Occupancy depends on the pool size *N* and the torque energy *ϵ*_*T*_, which increases with load but is not equal to it [24]. Occupancy is largest at high *ϵ*_*T*_ and large pool size (Fig. 4A). We defined mechanosensitivity as

**Figure 4.**
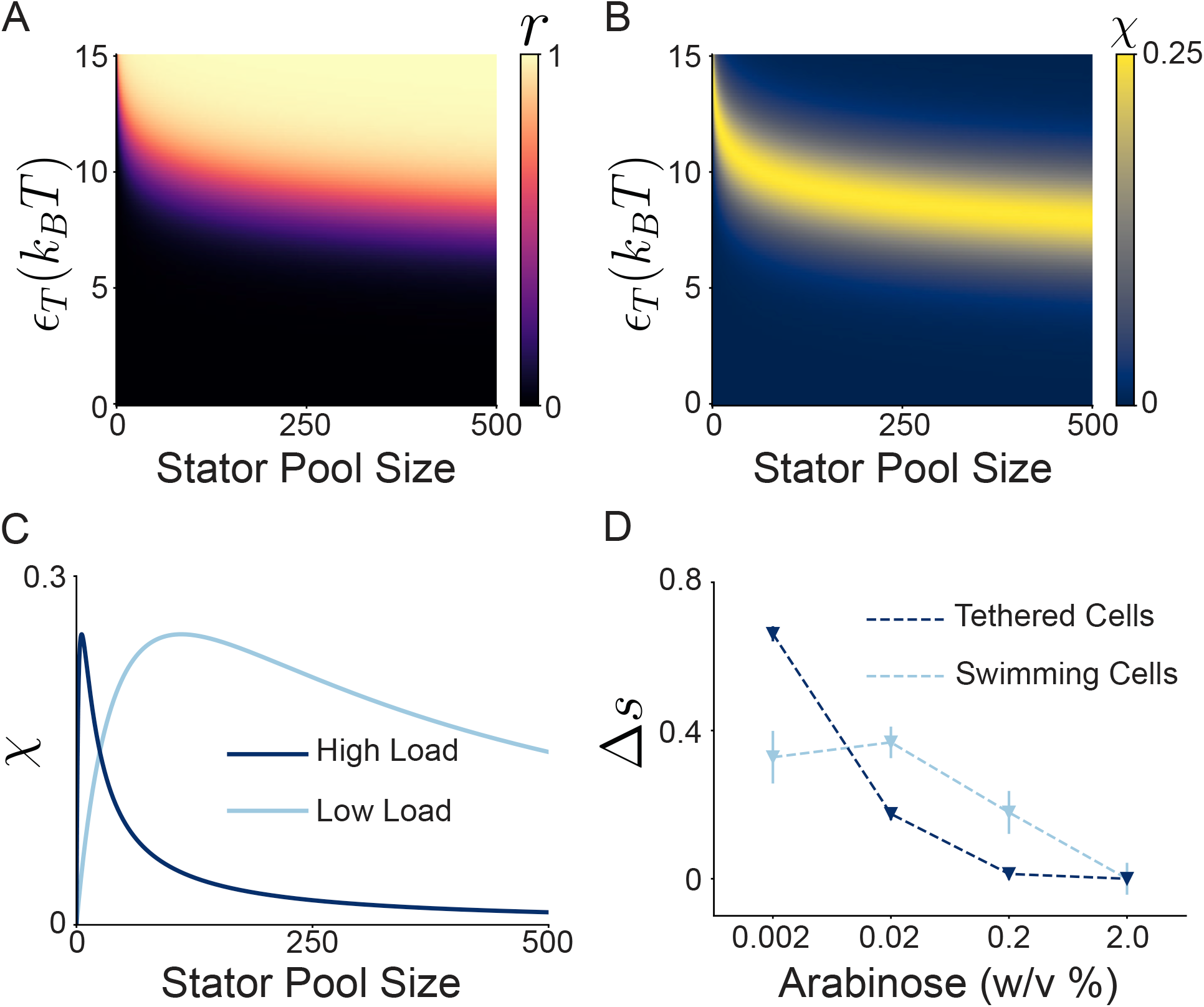
Stator pool size shapes mechanosensitivity of flagellar motors. **A)** A heatmap of average stator occupancy (*r*, unitless) against stator pool size and torque energy. Colorbar is shown to the right of the plot. **B)** A heatmap of mechanosensitivity (χ) against stator pool size and torque energy. Colorbar is shown to the right of the plot. **C)** Plots of mechanosensitivity (χ) against stator pool size for constant loads as predicted by our model. High load and low load are shown in dark and light blue respectively. **D)** The effective change in stator occupancy, Δ*s*, calculated from experiments. Δ*s* is a normalized change in speed between 0 and 15% Ficoll and is dimensionless. Error bars represent one standard error above and below the mean across biological replicates (counts in Tables S2 and S4).

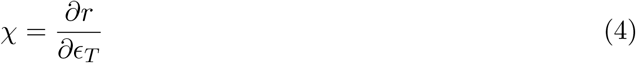

which represents how strongly occupancy responds to a change in load. Mechanosensitivity is high only within a restricted region of (*N, ϵ*_*T*_ ) space (Fig. 4B). At small pool sizes, motors are mechanosensitive only at high load, and larger pools shift the sensitive range to lower loads. At high load, only motors with small pools remain mechanosensitive (Fig. 4C).

To estimate χ experimentally, we normalized each speed by the speed at 2% arabinose in the same Ficoll concentration, where motors are saturated (speed did not rise above 2% arabinose; Fig. S4), so that the normalized speed reports fractional occupancy. The effective change in occupancy, Δ*s*, is the difference between this normalized speed at 15% and 0% Ficoll for a given arabinose concentration (see Supporting Material for details), a finite-difference analog of *χ*: a larger Δ*s* means the motor gained more occupancy as load increased. For swimming cells, Δ*s* was similar at the two lowest arabinose concentrations and fell to zero as expression increased. For tethered cells, Δ*s* dropped sharply at low arabinose and then plateaued (Fig. 4D). Because a tethered cell loads its motor more heavily than a swimming cell does, the difference between the assays follows from the model (Fig. 4C): higher load confines mechanosensitivity to smaller pools. The measured Δ*s* thus agrees with the prediction that pool size sets the load range over which the motor is mechanosensitive.

## Discussion

Stator pool size tunes the mechanosensitivity of the flagellar motor by setting the rate at which stators bind, which controls the load range over which the motor adapts. Our model predicts that a larger stator pool raises steady-state occupancy at low load but has no effect at high load, where the motor saturates (Fig. 1D). Experiments supported both parts of this prediction: swimming speed increased with stator expression in motility buffer, while at 15% Ficoll the differences among populations were no longer detectable (Fig. 2D). Single-motor measurements in tethered cells reproduced this load-dependent pattern (Fig. 3C), and because they load their motors more heavily than flagellar bundles do, they reached the load-insensitive regime at a lower Ficoll concentration than swimming cells. These trends follow from a balance of two rates: the on-rate *k*_+_ scales with the stator pool, while the off-rate *k*_−_ falls with load through the stator−peptidoglycan catch bond. At high load the off-rate is suppressed enough that the motor saturates regardless of pool size, whereas at low load occupancy is set by *k*_+_ and therefore by *N* . Together, these results identify the cellular stator pool as a determinant of motor mechanosensitivity, alongside the previously characterized catch-bond mechanism (Fig. 4).

The qualitative agreement between our model and the measured trends is notable because the model parameters were determined from prior measurements rather than tuned to our data (Table S1). However, one limitation of our study is that we do not know the exact size of the stator pool at each arabinose concentration. The pBAD33 system, although tightly regulated, can express heterogeneously across cells [42], making a direct calibration of pool size against arabinose concentration difficult. Even so, swimming speed rose monotonically with arabinose at 0% Ficoll (Fig. 2D). At low Reynolds number, speed reflects motor rotation rate and hence the number of stators driving the motor, so this steady rise points to more bound stators and, at this low load, to a larger pool. Swimming speed also depends on the number and arrangement of flagella and on bundle hydrodynamics, which we assume do not vary systematically with stator expression. The tethered-cell assay, which isolates a single motor, supports this assumption by reproducing the same load-dependent pool effect. Our western blots support this directly, showing a clear rise in stator expression from 0% to 0.02% arabinose, although the further increase is not resolved due to limits of the blot’s sensitivity (Figs. 2A, S1). Other studies using the same *motAB*-pBAD33 construct report comparable blots [43, 44]. We therefore treat the stator pool as increasing with arabinose, without assigning it an absolute value.

Our low-load result should not be confused with the zero-load limit. Near zero external load, motor speed is set by the rate of ion transit and is independent of stator number [34, 35]. A freely swimming cell, however, still works against the drag of its flagellar bundle, an intermediate load at which occupancy, and hence speed, rises with pool size. The pool dependence we observe at low load is therefore not contradicted by the zero-load result. We treat load throughout as a qualitative axis from low to high rather than a calibrated *ϵ*_*T*_, since neither Ficoll concentration nor tethering geometry maps directly onto the model’s torque energy. Because of its high molecular mass, Ficoll 400 increases viscosity with little change in medium osmolality. Although *E. coli* transiently responds to Ficoll as an attractant, clockwise bias returns toward baseline within about 2 min [46]. Cells remained in spatially uniform Ficoll for 5–10 min before imaging, making this response unlikely to explain the observed trends.

The wild-type pool of roughly 100 stators per cell likely reflects a trade-off between recruiting stators quickly and keeping the motor able to adapt. A larger pool shortens the diffusion-limited time for a stator to bind to the motor, raising the on-rate *k*_+_. But *k*_+_ must be matched to the off-rate *k*_−_, set by catch-bond kinetics, for the motor to stay mechanosensitive (Fig. 4). At the load of a flagellar bundle, a wild-type motor runs with only about half its full complement of stators, 5 to 6 of 11 [32], leaving room to recruit more as load rises while still swimming at a reasonable speed. A pool that is too small holds the motor below saturation across all loads, limiting torque; a pool that is too large saturates the motor even at low load and removes its ability to adapt.

Using a simple energetic argument, we speculate that stator pool size may be under evolutionary selection. A single stator contains 2091 amino acids (5 MotA and 2 MotB), and synthesizing one amino acid costs about 6 ATP [47], so building 100 stators consumes about 1.25 × 10^6^ ATP, roughly 0.0042% of a cell’s total budget of about 3 × 10^10^ ATP [47]. We treat this fractional cost as an upper-bound selection coefficient (*s* = 4.2 × 10^×5^); it is possible that the true value is lower due to motility benefits. Selection overcomes drift in a haploid population when 2*N*_*e*_*s >* 1, which requires *N*_*e*_ *>* 1.2 × 10^4^. The effective population size of *E. coli* lies between 10^5^ and 10^8^ [48], well above this threshold. The cost of maintaining the stator pool is therefore large enough to be under selection, suggesting that the observed pool size has been tuned to balance baseline performance against adaptive range.

Our two-state model is a deliberate simplification of stator dynamics, but the pool-size conclusions reported here are robust to elaboration. The stator pool *N* enters the model through the on-rate, *k*_+_ ∝ *N*, so it affects only the unbound-to-bound transition; once a stator is bound, transitions among bound substates do not depend on pool size. Two such substates have been characterized: a short-lived “hidden state,” thought to occur when a bound stator briefly detaches from the rotor and stops generating torque [33], and a split of the bound state into loosely and tightly bound configurations during torque generation [29]. Including them yields a four-state model that would sharpen predictions for torque-dependent dynamics but leave the pool-size effects unchanged. Our model also idealizes the diffusing state, treating stators as non-interacting with each other and with other membrane proteins. Membrane crowding, represented by a change in *γ*, should slow binding and shift mechanosensitivity, an effect we do not test here.

The principle we describe is not necessarily specific to the flagellar motor. Any force-responsive assembly that draws its components from a reversible pool could have its sensitive range of load set by the abundance of those components, just as stator pool size does for the motor here. The same logic may apply to focal adhesions and to cytoskeletal structures whose subunits recycle through a cytoplasmic pool [36, 37]. Component abundance then becomes a simple way to position a mechanosensor’s dynamic range, one that cells may tune by expression and that synthetic biology could exploit to build sensors with a chosen range of response.

## Data and code availability

Before publication, processed data and analysis code will be made publicly available at https://github.com/wadhwalab/2026-Wise-Stator-Pool, and raw video files will be deposited in a public data repository.

## Supporting material

Supporting material includes a model derivation, mechanosensitivity analysis, four figures, and six tables.

## Acknowledgments

We thank Seiga Yanagisawa for assistance with research design. This work was supported by the National Institute of General Medical Sciences of the National Institutes of Health (grant R00GM134124) and by the Arizona Biomedical Research Centre (grant RFGA2023-008-14) to N.W.

## Author Contributions

B.M.W. and N.W. designed research; B.M.W. derived model equations and performed simulations; S.A.S. generated strains and performed western blots; B.M.W. performed swimming and tethered-cell assays; B.M.W. analyzed data; all authors wrote the manuscript.

