## Supplemental Information for "Stator pool size shapes the mechanosensitivity of the bacterial flagellar motor"

### Supporting Information

#### Model

From Wadhwa et al (2019), we find the expression for the ratio of rate constants:

$$\frac{k_+}{k_-} = e^{-\frac{1}{k_B T}(\epsilon_b - \epsilon_T - \mu)}$$

Following previous studies, we assume that the load (torque) dependence belongs to $k_-$ through the catch-bond mechanism, meaning

$$k_+ \propto e^{-\frac{1}{k_B T}(-\mu)}$$

where the proportionality constant is $k_0$, as used in our Gillespie simulations. Chemical potential is defined as the free energy difference ($\Delta F$) contribution of a single molecule entering the system, where $F \equiv U - TS$. In the case of the motor, the chemical potential is the free energy difference between having $N$ and $N-1$ stators in the pool.

$$\mu = F(N) - F(N-1)$$

$$= N\epsilon_{\text{sol}} - k_B T \ln \frac{\Omega^N}{N!} - (N-1)\epsilon_{\text{sol}} + k_B T \ln \frac{\Omega^{N-1}}{(N-1)!}$$

where $\epsilon_{\text{sol}}$ is the energy of a stator in solution (in the pool) and $\Omega$ is the number of lattice sites. In a lattice model we assume that the solution can be divided into evenly sized areas (lattices) which can each hold a single stator. The particle number $N$ in this lattice gas is the stator pool size used in the main text, of order 100.

$$\mu = \epsilon_{\text{sol}} + k_B T \ln \frac{\Omega^{N-1}}{(N-1)!} \frac{N!}{\Omega^N} = \epsilon_{\text{sol}} + k_B T \ln \frac{\Omega^N \Omega^{-1}}{\frac{N!}{N}} \frac{N!}{\Omega^N}$$

$$\mu = \epsilon_{\text{sol}} + k_B T \ln \frac{N}{\Omega} \tag{S1}$$

The number of lattice sites can physically be understood as the surface area of the inner membrane divided by the area that a stator occupies, which we denote $\frac{1}{\gamma}$. Therefore, $\mu = \epsilon_{\text{sol}} + k_B T \ln N\gamma$. We can now rewrite the expression for the on-rate using properties of logarithms. From this point forward, we use $\beta = \frac{1}{k_B T}$ for simplicity.

$$k_+ \propto e^{\beta(\epsilon_{\text{sol}} + k_B T \ln N\gamma)} = e^{\beta\epsilon_{\text{sol}}} e^{\ln(N\gamma)}$$

The final expression for the on-rate is:

$$k_+ \propto N\gamma e^{\beta\epsilon_{sol}} \tag{S2}$$

The steady-state solution to equation 1 of the main text occurs when $\frac{d\bar{n}}{dt} = 0$

$$0 = k_+ n_{\text{max}} - k_+ n_{\text{ss}} - k_- n_{\text{ss}}$$

$$n_{\text{ss}}(k_+ + k_-) = k_+ n_{\text{max}}$$

$$n_{\text{ss}} = \frac{n_{\text{max}}}{1 + \frac{k_-}{k_+}} = \frac{n_{\text{max}}}{1 + \frac{e^{\beta(\epsilon_b - \epsilon_T)}}{N\gamma e^{\beta\epsilon_{\text{sol}}}}} \tag{S3}$$

The general solution to Eqn. (1) is:

$$\bar{n}(t) = n_{ss} + (n_0 - n_{ss})e^{-(k_+ + k_-)t}$$

where $n_0$ is the initial number of stators. This follows exponential decay with a time constant $\tau = \frac{1}{k_+ + k_-}$.

#### Mechanosensitivity

By rewriting Eqn. (S3) with $r = \frac{n_{\text{ss}}}{n_{\text{max}}}$, where $0 < r < 1$ is the occupancy, we find

$$r = \frac{1}{1 + \frac{e^{\beta(\epsilon_b - \epsilon_T)}}{N\gamma e^{\beta\epsilon_{\text{sol}}}}}$$

As described in the main text, $\chi = \frac{\partial r}{\partial \epsilon_T}$. We can find a closed form expression for $\chi$ as follows:

$$\chi = \frac{\partial r}{\partial \epsilon_T} = \frac{\partial r}{\partial e^{-\beta\epsilon_T}} \cdot \frac{\partial e^{-\beta\epsilon_T}}{\partial \epsilon_T}$$

$$\chi = \frac{N\gamma e^{\beta(\epsilon_b + \epsilon_{\text{sol}})}}{(e^{\beta(\epsilon_b - \epsilon_T)} + N\gamma e^{\beta\epsilon_{\text{sol}}})^2} \cdot \beta e^{-\beta\epsilon_T}$$

$$\chi = \frac{\beta N\gamma e^{\beta(\epsilon_b - \epsilon_T + \epsilon_{\text{sol}})}}{(e^{\beta(\epsilon_b - \epsilon_T)} + N\gamma e^{\beta\epsilon_{\text{sol}}})^2} \tag{S4}$$

$\chi$ has units of $[\text{Energy}]^{-1}$, which is consistent with our definition.

In order to understand mechanosensitivity from our experimental data we considered a change in speed (linear speed ($v$) and rotational frequency ($f$) for swimming and tethered-cell assays respectively) between 15 and 0% Ficoll. We cannot directly compare speeds at 15 and 0% Ficoll due to the change in viscosity, so we divided speed values by a reference speed. We chose our reference speed to be the speed of cells at 2% arabinose. A change in speed corresponds to a change in stator occupancy, so we calculated

$$\Delta s = \frac{v_k(15\%\ \text{Ficoll})}{v_{2\%}(15\%\ \text{Ficoll})} - \frac{v_k(0\%\ \text{Ficoll})}{v_{2\%}(0\%\ \text{Ficoll})} \tag{S5}$$

for each arabinose concentration, $k$. This is what we call the effective change in stator occupancy.

We also repeated this for our tethered-cell assay data, using the rotational frequency $f$ instead of linear speed $v$. By definition, $\Delta s$ for 2% arabinose is 0.

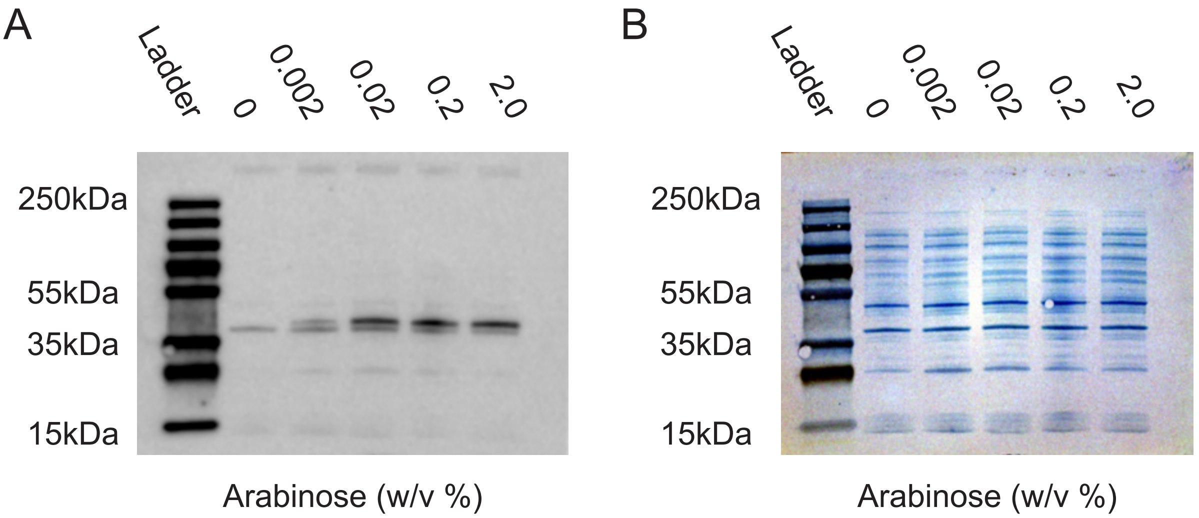


Supplemental Figure S1: **Western blot showing change in MotB. A)** Western blot of 6×His-tagged MotB under expression from pBAD33. Wells (left to right) are protein ladder, 0% arabinose, 0.002% arabinose, 0.02% arabinose, 0.2% arabinose, and 2% arabinose. **B)** Coomassie stain of Western blot. Wells (left to right) are protein ladder, 0% arabinose, 0.002% arabinose, 0.02% arabinose, 0.2% arabinose, and 2% arabinose.

Supplemental Table S1: **Parameters for simulations** $k_0$ is the timescale of stator binding, $\gamma$ is the ratio of a stator's cross-sectional area to the total inner-membrane area, $\epsilon_{\mathrm{sol}}$ is the unbound energy, $\epsilon_{\mathrm{b}}$ is the bound energy, and $\epsilon_{\mathrm{T}}$ is the torque energy.

| Parameters | Low Load | High Load | Source |
|---|---|---|---|
| $k_0$ ($\mathrm{s}^{-1}$) | 0.0015 | 0.0015 | Wadhwa et al. (2019), Tipping et al. (2013), Shi et al. (2019) |
| $\gamma$ | 0.0006 | 0.0006 | Santiveri et al. (2020) |
| $\epsilon_{\mathrm{sol}}$ ($k_BT$) | 3 | 3 | Estimate based on Wadhwa et al. (2019) |
| $\epsilon_b$ ($k_BT$) | 8 | 8 | Estimate based on Wadhwa et al. (2019) |
| $\epsilon_T$ ($k_BT$) | 9.5 | 12.5 | Estimate based on Wadhwa et al. (2019) |

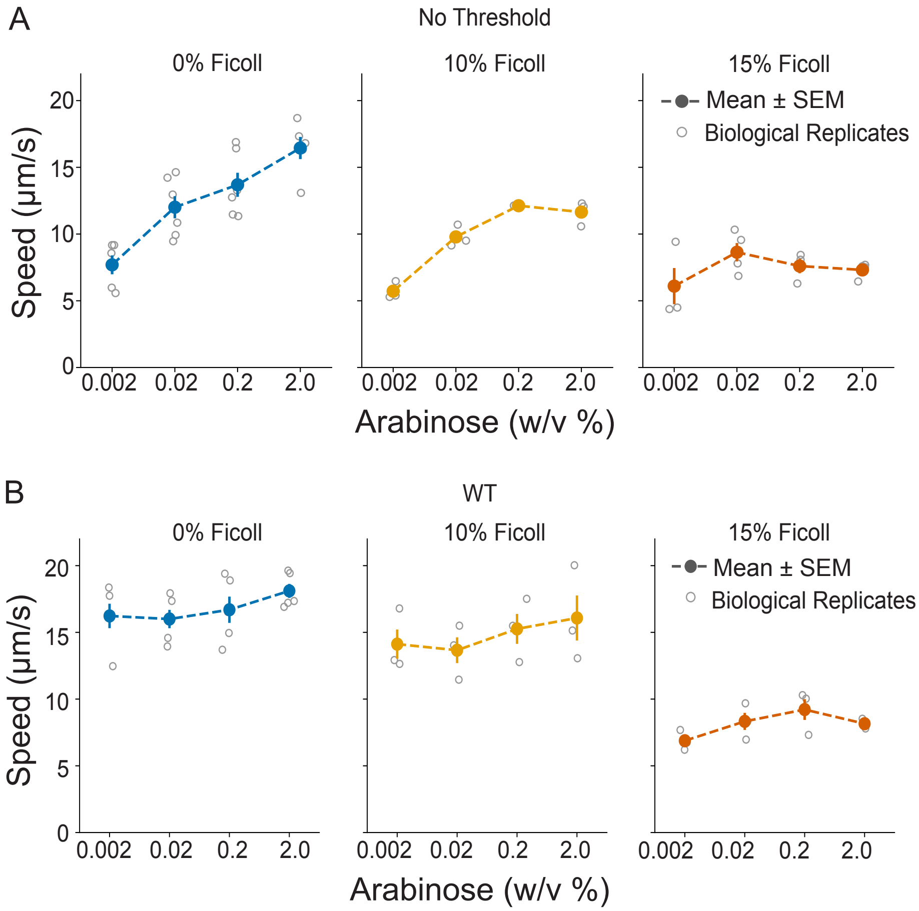


Supplemental Figure S2: **Control experiments for swimming assays.** **A)** Average swimming speeds of swimming cells at various stator expression levels in 0%, 10%, and 15% Ficoll (blue, yellow, and orange respectively) with no speed threshold applied. The error bars represent one standard error above and below the mean, and gray circles are the averages of individual biological replicates. **B)** Average swimming speeds of wild-type cells at various stator expression levels in 0%, 10%, and 15% Ficoll (blue, yellow, and orange respectively). The error bars represent one standard error above and below the mean, and gray circles are the averages of individual biological replicates.

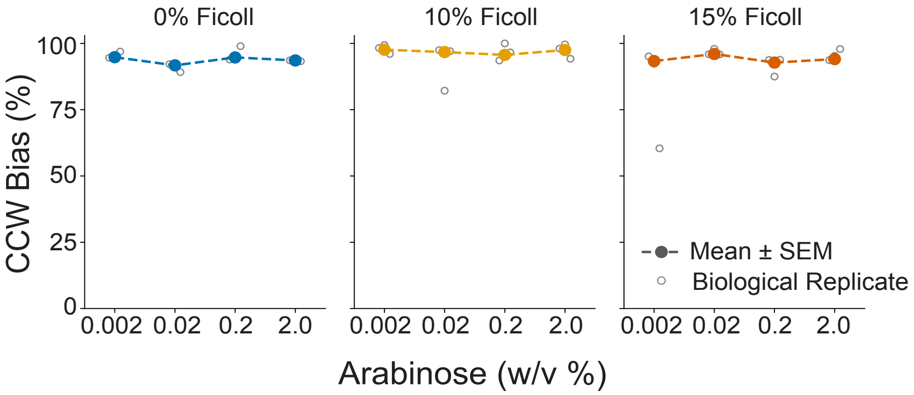


Supplemental Figure S3: **CCW bias did not change with stator expression and change in viscosity.** CCW bias is defined as the fraction of a recording in which a cell is rotating CCW (positive frequency). Average CCW bias of tethered cells at various stator expression levels in 0%, 10%, and 15% Ficoll (blue, yellow, and orange respectively). The error bars represent one standard error above and below the mean, and gray circles are the averages of individual biological replicates.

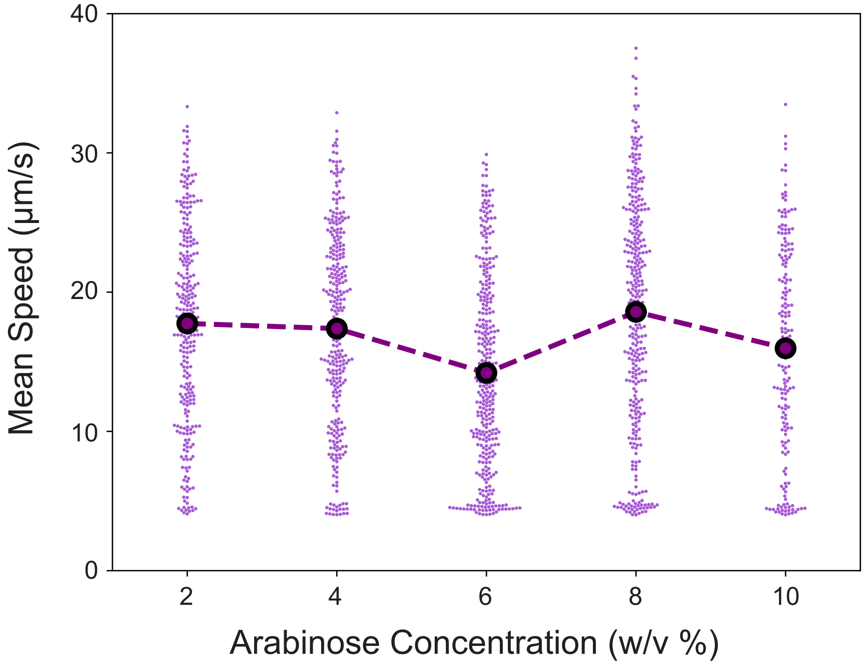


Supplemental Figure S4: **Speeds of cells above 2% arabinose.** Average swimming speeds of NW166 in MB + 1 mM methionine at increased arabinose concentration are shown by large markers. Smaller dots represent speeds of individual cells. The non-motile threshold has been applied.

Supplemental Table S2: **Summary of Swimming Assays** Number of motile cells analyzed for strain NW166 across all replicates for each condition.

| Arabinose (%) | Ficoll (%) | Motile Cells | Biological Replicates |
|---|---|---|---|
| 2 | 0 | 345 | 6 |
| 2 | 10 | 232 | 3 |
| 2 | 15 | 106 | 3 |
| 0.2 | 0 | 283 | 6 |
| 0.2 | 10 | 191 | 3 |
| 0.2 | 15 | 90 | 3 |
| 0.02 | 0 | 158 | 6 |
| 0.02 | 10 | 226 | 3 |
| 0.02 | 15 | 124 | 3 |
| 0.002 | 0 | 373 | 6 |
| 0.002 | 10 | 204 | 3 |
| 0.002 | 15 | 108 | 3 |

Supplemental Table S3: **Summary of Swimming Assays** Number of motile cells analyzed for strain MG1655 across all replicates for each condition.

| Arabinose (%) | Ficoll (%) | Motile Cells | Biological Replicates |
|---|---|---|---|
| 2 | 0 | 989 | 5 |
| 2 | 10 | 592 | 3 |
| 2 | 15 | 249 | 3 |
| 0.2 | 0 | 817 | 5 |
| 0.2 | 10 | 814 | 3 |
| 0.2 | 15 | 629 | 3 |
| 0.02 | 0 | 748 | 5 |
| 0.02 | 10 | 526 | 3 |
| 0.02 | 15 | 333 | 3 |
| 0.002 | 0 | 522 | 5 |
| 0.002 | 10 | 420 | 3 |
| 0.002 | 15 | 224 | 3 |

Supplemental Table S4: **Summary of Tethered-Cell Assays** Number of motile cells analyzed for strain NW263 across all replicates for each condition.

| Arabinose (%) | Ficoll (%) | Number of Cells | Biological Replicates |
|---|---|---|---|
| 2 | 0 | 104 | 3 |
| 2 | 10 | 31 | 3 |
| 2 | 15 | 48 | 3 |
| 0.2 | 0 | 64 | 3 |
| 0.2 | 10 | 35 | 3 |
| 0.2 | 15 | 31 | 3 |
| 0.02 | 0 | 113 | 3 |
| 0.02 | 10 | 50 | 3 |
| 0.02 | 15 | 34 | 3 |
| 0.002 | 0 | 51 | 3 |
| 0.002 | 10 | 30 | 3 |
| 0.002 | 15 | 23 | 3 |

Supplemental Table S5: **Summary of ANOVA Results** For each experiment (swimming assay or tethered-cell) we report the p-values from ANOVA between the replicate means of all four groups (arabinose concentrations).

| Experiment | Ficoll (%) | p-value |
|---|---|---|
| Swimming | 0 | $< 0.001$ |
| Swimming | 10 | $< 0.001$ |
| Swimming | 15 | 0.478 |
| Swimming (WT) | 0 | 0.371 |
| Swimming (WT) | 10 | 0.684 |
| Swimming (WT) | 15 | 0.176 |
| Tethered | 0 | $< 0.001$ |
| Tethered | 10 | 0.149 |
| Tethered | 15 | 0.091 |

Supplemental Table S6: **Summary of TOST Results** For each experiment (swimming assay or tethered-cell) we performed a TOST test on the replicate means. This table shows $\Delta = \mu_2 - \mu_1$, the 90% confidence interval for $\Delta$, and the p-value. Equivalence intervals were pre-defined as $\pm 3\mu$m/s for swimming assays and $\pm 1.3$Hz for tethered-cell assays. $\Delta$ and the confidence interval are in units of $\mu$m/s for swimming assays and Hz for tethered-cell assays.

| Experiment | Ficoll (%) | Arabinose 1 (%) | Arabinose 2 (%) | $\Delta$ | 90% C.I. | p-value |
|---|---|---|---|---|---|---|
| Swimming | 10 | 2 | 0.2 | 0.538 | (-0.917, 1.994) | 0.017 |
| Swimming | 15 | 2 | 0.2 | 0.019 | (-3.585, 3.624) | 0.075 |
| Swimming | 15 | 0.2 | 0.02 | 0.907 | (-2.725, 4.539) | 0.133 |
| Swimming | 15 | 0.02 | 0.002 | 2.397 | (-1.19, 5.983) | 0.358 |
| Swimming | 15 | 2 | 0.02 | -0.888 | (-3.063, 1.287) | 0.054 |
| Swimming | 15 | 2 | 0.002 | 1.509 | (-2.056, 5.074) | 0.208 |
| Swimming | 15 | 0.2 | 0.002 | 1.490 | (-2.632, 5.612) | 0.239 |
| Tethered | 10 | 2 | 0.2 | 1.944 | (-1.021, 4.908) | 0.683 |
| Tethered | 10 | 2 | 0.02 | 1.725 | (-1.257, 4.706) | 0.626 |
| Tethered | 10 | 2 | 0.002 | 2.331 | (-0.856, 5.518) | 0.775 |
| Tethered | 10 | 0.02 | 0.002 | 0.606 | (-0.582, 1.795) | 0.124 |
| Tethered | 10 | 0.002 | 0.2 | 0.388 | (-0.898, 1.673) | 0.091 |
| Tethered | 10 | 0.02 | 0.2 | 0.219 | (-1.145, 1.583) | 0.083 |
| Tethered | 15 | 2 | 0.2 | 0.445 | (-0.185, 1.074) | 0.023 |
| Tethered | 15 | 2 | 0.02 | -0.019 | (-1.646, 1.607) | 0.078 |
| Tethered | 15 | 2 | 0.002 | 0.814 | (-1.394, 3.021) | 0.305 |
| Tethered | 15 | 0.02 | 0.002 | 0.833 | (-1.346, 3.012) | 0.333 |
| Tethered | 15 | 0.002 | 0.2 | 0.369 | (-1.879, 2.617) | 0.182 |
| Tethered | 15 | 0.02 | 0.2 | 0.464 | (-1.199, 2.127) | 0.149 |
